# The TMJ disc is a common ancestral feature in all mammals, as evidenced by the presence of a rudimentary disc during monotreme development

**DOI:** 10.1101/2020.01.17.910471

**Authors:** Neal Anthwal, Abigail S Tucker

## Abstract

The novel mammalian jaw joint, known in humans as the temporomandibular joint or TMJ, is cushioned by a fibrocartilage disc. This disc is secondarily absent in therian mammals that have lost their dentition, such as giant anteaters and some baleen whales. The disc is also absent in all monotremes. However, it is not known if the absence in monotremes is secondary to the loss of dentition, or if it is an ancestral absence. We use museum held platypus and echidna histological sections to demonstrate that the developing monotreme jaw joint forms a disc primordium that fails to mature and become separated from the mandibular condyle. We then show that monotreme developmental anatomy is similar to that observed in transgenic mouse mutants with reduced musculature. We therefore suggest that the absence of the disc on monotremes is a consequence of the changes in jaw musculature associated with the loss of adult teeth. Taken together, these data indicate that the ancestors of extant monotremes likely had a jaw joint disc, and that the disc evolved in the last common ancestor or all mammals.

## Introduction

The temporomandibular joint (TMJ) is the one of the most used joints in the body, articulating the upper and lower jaw in mammals. A fibrous articular disc sits between the skeletal elements of the joint and acts as a cushion.

TMJ development occurs by the coming together of two membranous bones: the condylar process of the dentary bone in the mandible and the squamosal bone in the skull. The interaction of the condylar with the squamosal induces the formation of a glenoid (or mandibular) fossa on the latter (Wang et al., 2011). The articular disc sits between the two within a synovial capsule. The TMJ disc attaches to the superior head of the lateral pterygoid muscle anteriorly, and to ligaments posteriorly including the disco-mallear ligament that runs thought the capsule of the middle ear, joining the malleus to the TMJ disc. The TMJ articulates the jaw in all mammals, and is referred to as the squamosal dentary joint (SDJ) in those mammals without a fused temporal bone. In non-mammals the upper and lower jaw articulate via the endochondral quadrate and articular, known as the primary jaw joint (Wilson and Tucker, 2004). TMJ developmental anatomy reflects its evolutionary history as this novel, jaw joint forms after the development of the primary joint, which, in mammals, is integrated into the middle ear (Takechi and Kuratani, 2010; Anthwal et al., 2013; Maier and Ruf, 2016; Tucker, 2017). In recent years, a number of studies have advanced the understanding of middle ear evolution in the context of anatomical development (Luo, 2011; Anthwal et al., 2013, 2017; Urban et al., 2017; Wang et al., 2019), but little work has sought to understand the TMJ in an evolutionary and comparative developmental biology context. This is despite the crucial role that the formation of the TMJ has in mammalian evolution.

An important part of the TMJ is the disc that cushions its action. The origin of the disc is uncertain. The insertion of the lateral pterygoid muscle into the disc on the medial aspect, and the presence of the discomalleolar ligament has led to speculation that the disc represents a fibrocartilage sesamoid with a tendon of a muscle of mastication trapped by the novel mammalian jaw joint (Herring, 2003). However, studies in mice indicate that the disc develops from a region of flattered mesenchyme cells adjacent to, or possibly part of, the perichondrium of the developing condylar cartilage (Purcell et al., 2009, 2012; Hinton et al., 2015). Formation of the disc condensation is dependent on Ihh signalling from the cartilage (Shibukawa et al., 2007; Purcell et al., 2009; Yang et al., 2016), and Fgf signalling via Spry 1 and 2 genes from the adjacent muscles (Purcell et al., 2012). Therefore, the disc may have its origins in either a tendon, the novel secondary cartilage of the condylar process, or a combination of the two.

Interestingly the disc is absent in extant monotremes (Sprinz, 1964). Monotremes and therian mammals (marsupials and placentals) are evolutionary distant, with the common ancestor of the two subclasses being a mammal like reptile form around 160 million years ago (Kemp, 2005). Monotremes have a number of “reptile” like anatomical features such as a cloaca, external embryonic development in an egg, a straight cochlear in the inner ear and laterally protruding legs (Griffiths, 1978). The absence of a disc in both echidna and platypus suggests that the disc evolved after the split between monotremes and therian mammals, and is therefore a therian novelty. Alternatively, absence of the TMJ disc in extant monotremes might be due to a secondary loss of this structure, linked to changes in mastication with the reduction and loss of teeth. Extant adult monotremes are edentulous, possibly due to the evolution of electroreceptivity (Asahara et al., 2016). The juvenile platypus has rudimentary teeth that regress (Green, 1937), while the echidna shows only thickening of the dental epithelium during development. In contrast, fossil monotremes have a mammalian tribosphenic dentition and were capable of chewing (Kemp, 2005). The presence or absence of a disc in such fossils is difficult to ascertain due to lack of preservation of soft tissue. In support of mastication playing a role in disc formation edentulous therian mammals, or those lacking enamel, often lack a disc. These species include some (but not all) baleen whales (El Adli and Deméré, 2015), giant ant eaters and sloths (Naples, 1999).

In order to address this uncertainty, we have examined the development of the TMJ in monotremes and made comparison with mouse developmental models where muscle development is perturbed.

## Materials and Methods

Platypus *(Ornithorhynchus anatinus)* and short-beaked echidna *(Tachyglossus aculeatus*) slides were imaged from the collections at the Cambridge University Museum of Zoology. Details of samples imaged are in table 1. All museum samples have been studied in previously published works (Watson, 1916; Green, 1937; Presley and Steel, 1978). Stages for platypus are after Ashwell (Ashwell, 2012). Staging of echidna H.SP EC5 is estimated by cross-referencing previous studies (Griffiths, 1978; Rismiller and McKelvey, 2003). CT cans of adult platypus were a gift of Anjali Goswami, the Natural History Museum, London. *Mesp1Cre;Tbx1flox (Tbx1CKO)* mice were derived as previously described (Anthwal et al., 2015).

**Table 1:** Museum held specimens used in the current study. CRL – Crown rump length. ^*^Estimate based on Ashwell, 2012. ^†^ Estimate based on Griffiths, 1978 and Rismiller & McKelvey, 2003.

| Species | Collection | ID | Estimated Stage | CRL |
| --- | --- | --- | --- | --- |
| *Ornithorhynchus anatinus* | Cambridge | Specimen X | P6.5* | 33mm |
| *Ornithorhynchus anatinus* | Cambridge | Specimen Delta | P10* | 80mm |
| *Ornithorhynchus anatinus* | Cambridge | Specimen Beta | P80* | 250mm |
| *Tachyglossus aculeatus* | Cambridge | Echidna H.SP EC5 | P18† | 83mm |

Tissue processing and histological staining: embryonic samples for histological sectioning were fixed overnight at 4 °C in 4 % paraformaldehyde (PFA), before being dehydrated through a graded series of ethanol and stored at −20°C. For tissue processing, samples were cleared with Histoclear II, before wax infiltration with paraffin wax at 60°C. Wax embedded samples were microtome sectioned at 8 µm thickness, then mounted in parallel series on charged slides.

For histological examination of bone and cartilage, the slides were then stained with picrosirius red and alcian blue trichrome stain using standard techniques.

## Results

If the TMJ disc is a therian novelty then no evidence of a disc would be expected in extant monotremes during development of the TMJ. The development of the jaw joint was therefore examined in museum held histological sections of developing post-hatching platypus and compared with the mouse.

As other authors have previously described (Purcell et al., 2009; Hinton, 2014), in the embryonic day (E) 16.5 mouse, the disc anlage is observed as thickened later of mesenchyme connected to the superior aspect of the condylar cartilage (Figure 1A). At postnatal day (P) 0, the disc has separated from the condylar and sits within the synovial cavity of the jaw joint (Figure 1B). In a platypus sample estimated to be 6.5 days post hatching, the TMJ had been initiated, but the joint cavity had not yet formed (Figure 1C,D). Close examination of the superior surface of the condylar cartilage revealed a double layer of thickened mesenchyme in the future fibrocartilage layer of the condylar (Figure 1 C’,D’). The outer layer is similar to that known to develop into the articular disc in therian mammals (Purcell et al., 2009). This thickened mesenchyme persisted in older samples, estimated to be P10, where the synovial cavity of the TMJ was beginning to form above (Figure 1 E,E’). In the most mature platypus sample examined (around P80) the fibrocartilage layer of the condylar process was thick and had a double-layered structure (Figure 1F). The outer layer was connected via a tendon to the lateral pterygoid muscle. At this late stage of postnatal development, the platypus puggle would have been expected to start leaving the burrow and to be eating a mixed diet, although full weaning does not occur until around 205 days post hatching (Rismiller and McKelvey, 2003). In the mature platypus, the condylar process sits within a glenoid fossa (Figure 1 F,G), which was not fully formed at earlier stages. A disc-like structure lying over the condylar and connected to the adjacent muscles was therefore evident in the platypus postnatally but did not lift off the condylar at any stage.

**Figure 1.**
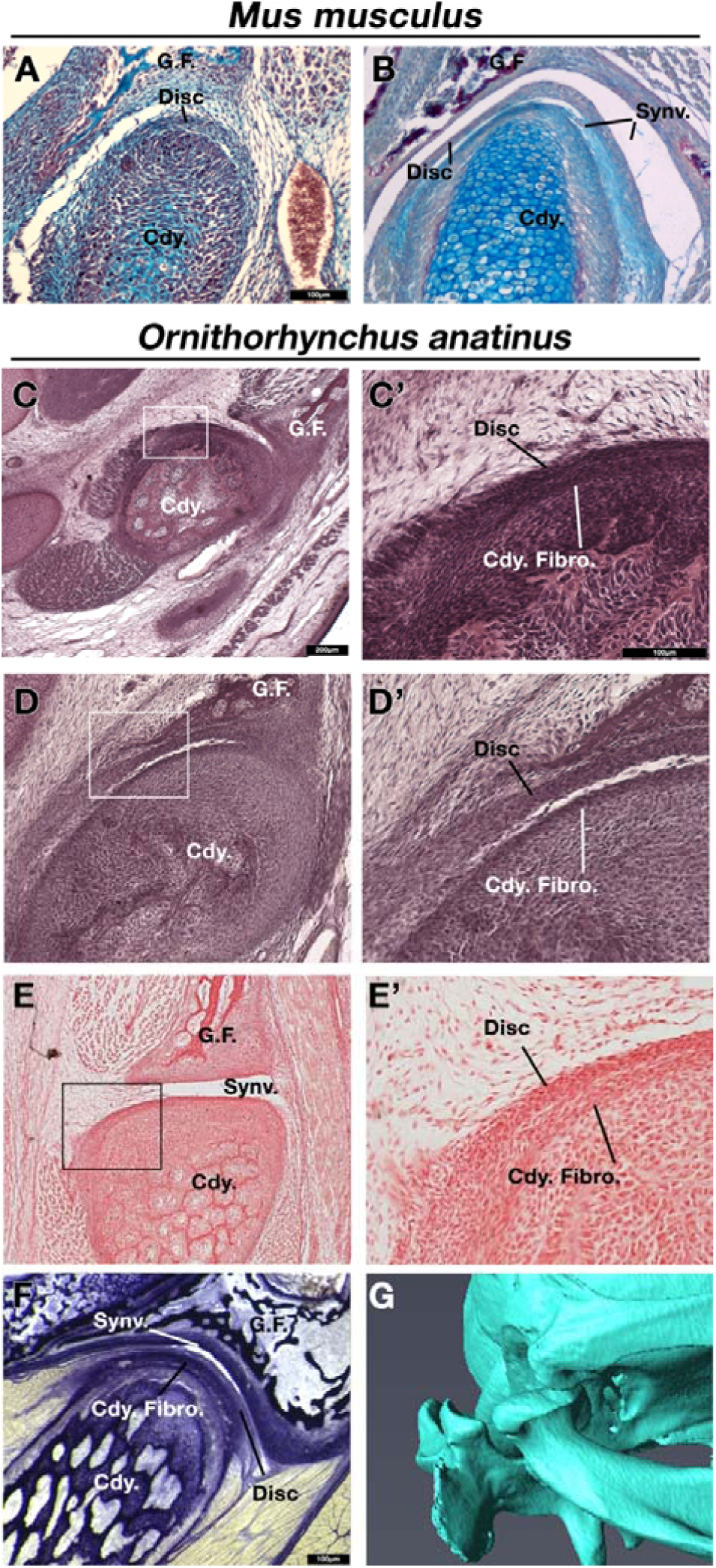
Comparison of mouse (Mus musculus) and platypus (Ornithorhynchus anatinus) developing jaw joint reveals the presence of a jaw joint disc anlage in early post-hatching platypus despite absence of the disc in adults. A,B Histological sections of mouse jaw joint disc development at embryonic day 16.5 (A) and postnatal day 0 (B).C-D’ Histological sections of estimated post hatching day 6.5 jaw joint at two different levels (C and D) Note that the separation between the disc anlage and condylar in D is probably a processing artefact. E,E’’ Histological sections of estimated post hatching day 10 jaw joint. F Histological section of mature jaw joint in a juvenile platypus are estimated post hatching day 80. G µCT scan of jaw joint region of adult platypus. G.F. – glenoid fossa; Cdy. – condylar process; Cdy. Fibro. – condylar fibrocartilage; Synv. – synovial cavity of the jaw joint.

Next we examined the development of the TMJ in a derived young short-beaked echidna puggle specimen with a crown-rump length of 83mm, which we estimate to be around P18. The TMJ is not fully (Figure 2). The condylar process possessed a thick, doubled fibrocartilage outer layer (Figure 2), much as was observed in the platypus (Figure 1D). The outer fibrocartilage later was connected by connective tissue to the lateral pterygoid muscle (Figure 2B’). Clear disc-like structures were therefore present during development in both extant monotremes.

**Figure 2.**
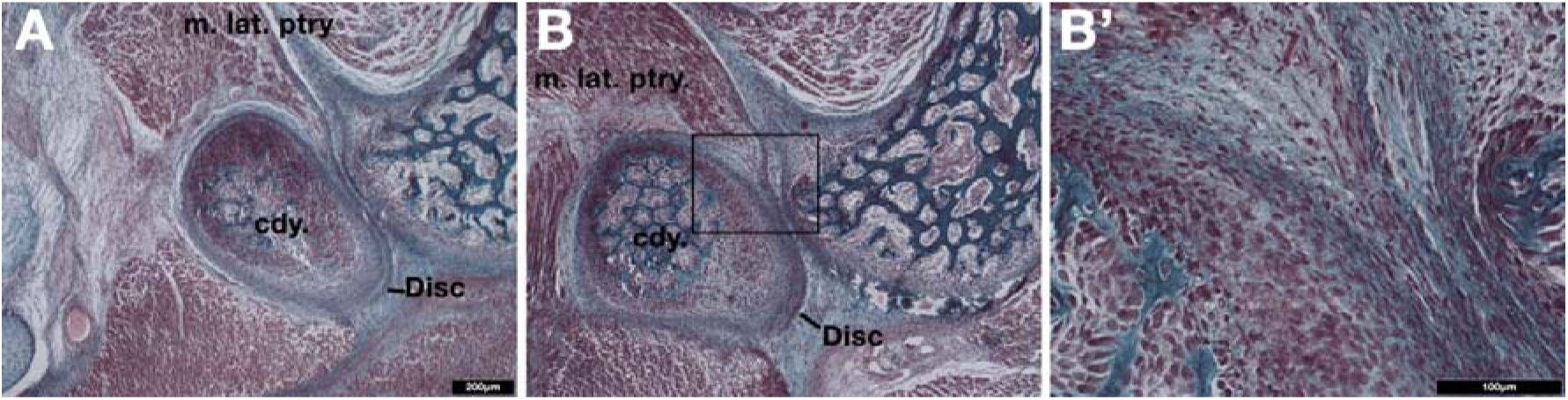
Examination of the developing jaw joint reveals the presence of a jaw joint disc anlage in post-hatching day 18 short-beaked echidna (Tachyglossus aculeatus). A-B Histological staining at the forming jaw articulation in echidna young estimated to be 18 days post hatching at two different level. Fibrocartilage disc anlage superior to the condylar and connected by tendon lateral pterygoid muscle is observed. B’ High-powered view of boxed region in B showing the connection between the muscle and the developing disc. Cdy. – condylar process; m. lat. ptry. – lateral pterygoid muscle.

Taken together, the developmental evidence suggests that extant monotremes initiate a layer of fibrocartilage connected to the lateral pterygoid muscle, similar to the initiation of the TMJ disc in therian mammals. However, unlike in therian mammals, the monotreme fibrocartilage failed to separate from the condylar to form an articular disc in the TMJ. Interactions with musculature, both mechanical (Habib et al., 2007; Purcell et al., 2012; Jahan et al., 2014; Nickel et al., 2018) and molecular (Shibukawa et al., 2007; Gu et al., 2008; Purcell et al., 2009, 2012; Kinumatsu et al., 2011; Michikami et al., 2012; Yasuda et al., 2012; Kubiak et al., 2016), have been suggested to be responsible for the proper formation of the TMJ disc. Lack of mechanical force in monotremes might therefore result in the disc remaining attached to the condylar. In order to examine how changes in mechanical loading affect disc development, we next examined disc development in the *Mesp1Cre;Tbx1flox* conditional mutant mouse (*Tbx1CKO*). This mouse has a mesoderm specific deletion of the T-box transcription factor *Tbx1*, resulting in hypomorphic muscle development (Grifone et al., 2008; Aggarwal et al., 2010; Anthwal et al., 2015).

We used alcian blue / alizarin red stained histological sections to investigate the development of the TMJ disc in *TbxCKO* mice at embryonic day 15.5. This is the stage when future disc mesenchyme is first observed (see Figure 1A). In wildtype embryos, the future disc mesenchyme was observed as a condensation attached to the superior surface of the condylar fibrocartilage (Figure 3A). A distinct disc-like mesenchyme was also observed superior to the condylar of the *Tbx1CKO* (Figure 3B). This mesenchyme and the fibrocartilage layer of the condylar cartilage both appeared thicker in the *Tbx1CKO* compared to its wildtype littermate. At E18.5, the wildtype TMJ disc had separated from the condylar process, and sat within a synovial joint cavity (Figure 3C). In the *Tbx1CKO* an upper synovial cavity had formed, similar to the WT, but the disc had failed to separate from the condylar (Figure 3D). Instead, a thickened band of fibrocartilage was observed on the superior surface of the condylar process.

**Figure 3.**
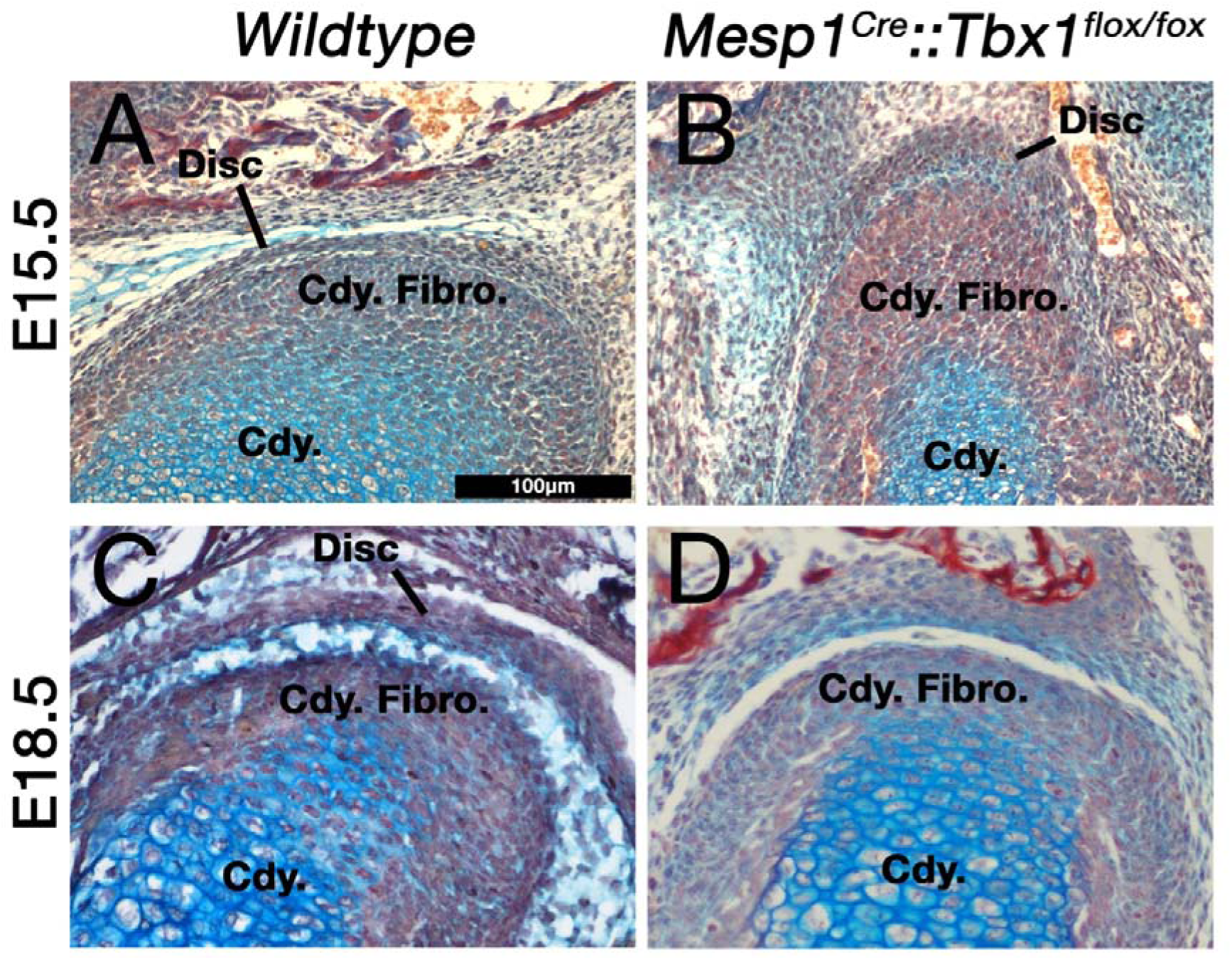
Muscle-disc interactions are required for the maturation and separation of the jaw joint articular disc. A,B The disc anlage is observed at E15.5 in both wildtype mice (A) and Mesp1Cre;Tbx1fl/fl mice with a hypomorphic muscle phenotype (B). C,D By E18.5 the disc has separated from the condylar process in wildtype mice (C), but not in and Mesp1Cre;Tbx1fl/fl mice. Cdy. – condylar process; Cdy. Fibro. – condylar fibrocartilage

## Discussion

The absence of an articular disc in monotremes has been thought to be either a secondary loss related to the absence of mature dentition, or the disc being a later acquisition in the therian clade. The data presented here show that a mesenchyme similar to the TMJ disc is initiated in both platypus and echidna jaws during post-hatching development, but fails to mature and separated from the dentary condyle. In the light of the failure of the disc to fully separate in transgenic mouse models with hypomorphic muscle development, it seems likely that the disc has been secondarily lost in edentulous mammals, including monotremes.

The secondary jaw joint of some of the earliest mammals-like reptiles with a mandibular middle ear, such as Morganuconodon, were able to withstand the biomechanical stresses sufficient for feeding on the hard keratinised bodies of insects, while others such as Kuehneotherium could not (Gill et al., 2014). Later animals developed a range of mandibular movements during chewing, including rolling, yaw and front to back movements (Kemp, 2005; Luo et al., 2015; Grossnickle, 2017; Lautenschlager et al., 2017, 2018; Bhullar et al., 2019). It Is not clear if these species had evolved an articular disc, since fibrocartilage is rarely fossilised. Based on the presence of the first stages of disc formation during monotreme development it is likely that the common stem Jurassic mammal-like reptilian ancestor of both monotremes and therian mammals had a disc. The data presented here confirms an essential biomechanical component in disc development. Therefore, we are able to consider when during mammalian evolution these forces were able to act to enable disc formation. For example, it is probable that many late Triassic and early Jurassic mammaliaforms such a Hadrocodium (Luo, 2001) possessed an articular disc, since they possessed a well formed squamosal dentary joint and occluding teeth capable of chewing.

One hypothesis for the origin of the articular disc is that it formed from the tendon of a muscle of jaw closure of the primary jaw joint interrupted by the formation of the novel mammalian jaw joint (Herring, 2003). The tendons and skeleton of the front of the head are derived from the cranial neural crest, whereas much of the musculature is mesoderm derived (Santagati and Rijli, 2003; Yoshida et al., 2008). Interactions between the mesoderm and neural crest co-ordinate the muscular skeletal development of the head (Grenier et al., 2009). A striking piece of evidence for the tendon origin of the disc is the expression in the developing articular disc of *Scleraxis* (Purcell et al., 2012; Roberts et al., 2019), a specific regulator of tendon and ligament development (Schweitzer et al., 2001; Sugimoto et al., 2013). If the disc is derived from a tendon, then it may be thought of as a fibrocartilage sesamoid. Such sesamoids are found in joints and in tendons that are subject to compression, like the tendons that pass around bony pulleys such as the flexor digitorum profundus tendon in quadrupeds, the patella tendon and ligament (Benjamin and Ralphs, 1998), and the cartilago transiliens in crocodilians (Tsai and Holliday, 2011). Fibrocartilages also form at the enthesis of long bones. Interestingly, it has been demonstrated that much like the TMJ disc, enthesis fibrocartilage cells are derived from Hh responsive cells and that these cells are responsive to mechanical loading (Schwartz et al., 2015). To support the tendon origin of the TMJ disc, our data show that the formation of the disc is dependent on interactions between the skeletal and muscle components of the TMJ. Such tissue interaction is also a key process in the formation of tendons and ligaments (Eloy-Trinquet et al., 2009; Huang, 2017).

The mechanism by which the disc fails to separate from the condylar in monotremes is not yet clear. Hh signalling is known to be involved in both the initiation of the disc, and the later separation from the condylar (Purcell et al., 2009). It is still possible that the role in Hh in separation of the disc is a therian innovation, and as such the reason that monotremes fail to do so is a lack of the later Hh dependent developmental programme for disc separation. However, the absence of the disc in therian edentates strongly suggests that the loss is secondary. Furthermore, the failure of the disc to elevate off the condylar in *Tbx1CKO*, with hypomorphic cranial musculature, suggests that the loss of discs in edentulous mammals is due to changes in the developmental biomechanics of the muscle/bone interactions that occurred as a consequence of loss of teeth, such as a reduction in size and power of the muscles of mastication. The formation and maturation of the disc is unlikely to be directly dependent on the presence of teeth. The TMJ disc forms normally during embryonic development in mice quite some time before the eruption of the teeth during the third postnatal week, while baleen whales vary in the presence or absence of TMJ discs, and indeed TMJ synovial cavities (El Adli and Deméré, 2015). In addition, it is clear that movement of the jaw is essential for maturation of the disc (see also (Habib et al., 2007)). Unfortunately, due to the rarity of fresh material, it is not possible to further examine the mechanistic aspects of TMJ development in edentulous monotreme species at the present time.

In conclusion, we demonstrate that during development, monotremes show evidence of initiation of the fibrocartilage articular disc, despite all adult monotremes not having an articular TMJ disc. The maturation and separation of the disc is dependent on biomechanical interactions with the associated musculature, as demonstrated by the failure of disc maturation and separation in mice mutants with hypomorphic cranial muscle. Therefore, toothed ancestors of monotremes likely had a TMJ disc. Our research suggests that changes in the cranial musculature that occurred as a consequence of a move towards edentulous dietary niches resulted in absence of the TMJ in monotremes, a parallel loss occurring in edentulous therian mammals (Figure 4). Finally, the presence of the disc anlage in monotremes indicates that the mammal-like reptile ancestors of all modern mammals likely possessed a disc to cushion the novel jaw articulation.

**Figure 4.**
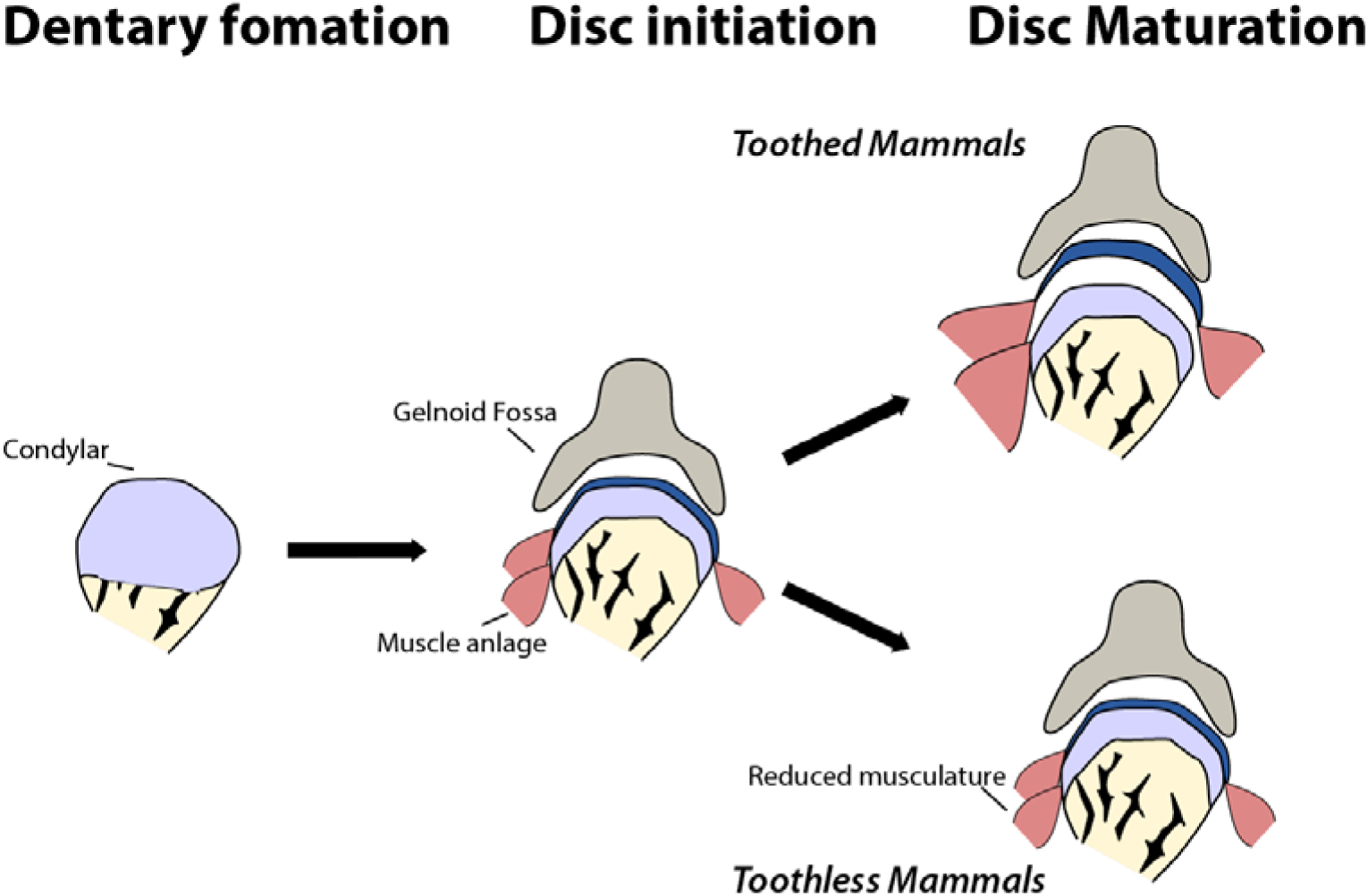
Maturation of the jaw joint articular disc in mammals is dependent on muscle interactions. In toothless mammals, reduction in jaw musculature results in changes in muscle-disc interaction and so the disc does not separate from the mandibular condyle to sit within the synovial joint capsule.

## Acknowledgments

We would like to thank and acknowledge the following people. Anjali Goswami provided µCT images of the adult platypus. Robert Asher provided access to samples held at the Zoological Museum in Cambridge University. Peter Giere provided access to the Hill Collection at the Berlin Museum für Naturkunde. Andrew Gillis provided assistance with imaging.

